# Causal evidence for LTP-based interference in visual learning

**DOI:** 10.1101/438598

**Authors:** Ji Won Bang, Diana Milton, Yuka Sasaki, Takeo Watanabe, Dobromir Rahnev

## Abstract

Training related skills in close succession results in interference but the reasons for this interference are not understood. Here we test the hypothesis that interference occurs due to competition of long-term potentiation (LTP): the LTP induced by one task impedes the LTP induced by the other. Human subjects performed two consecutive training sessions on different Gabor orientations. Immediately after the offset of the first training, we applied continuous theta burst stimulation (cTBS) to interfere with the LTP processes produced by the first training. We found that cTBS to a control site (vertex) resulted in substantial anterograde interference for the second training. Critically, cTBS to the visual cortex not only disrupted learning on the immediately preceding training, but also released the second training from the anterograde interference. These results provide strong support for the LTP-based theory of interference and suggest the possibility of directly manipulating the competition between different learning periods.

## Introduction

Training two related skills in close succession results in interference. Two types of interference have been observed: anterograde interference (i.e., new learning disrupting future learning) and retrograde interference (i.e., new learning disrupting a previous learning). Both types of interference are commonly observed across motor, perceptual, and cognitive learning (Cantarero et al. 2013; Leow et al. 2013; Wigmore, Tong, and Flanagan 2002; Sing and Smith 2010; Brashers-Krug, Shadmehr, and Bizzi 1996; Shadmehr and Brashers-Krug 1997; Seitz et al. 2005; Yotsumoto, Chang, et al. 2009; Shibata et al. 2017).

It is theorized that such interference is due to competition in long-term potentiation (LTP) processes (Cantarero et al. 2013; Ziemann et al. 2004; Stefan et al. 2006) since LTP is known to be a critical component of consolidating new learning (Beste et al. 2011; Nicoll 2017; Lynch 2004; Sanes and Donoghue 2000). Specifically, LTP induced by one learning is thought to interfere with LTP induced by another learning thus resulting in retrograde and anterograde interference.

Indirect evidence for this LTP-based theory of interference has come from research employing transcranial magnetic stimulation (TMS) to disrupt the post-training neural processes. Such studies have demonstrated that TMS to the primary motor cortex after motor training (Baraduc et al. 2004; Muellbacher et al. 2002; Robertson, Press, and Pascual-Leone 2005) or to the early visual cortex after visual training (De Weerd et al. 2012) can disrupt learning. However, these previous studies only employed learning of a single stimulus and therefore it remains unclear what the effects of TMS would be on the interference between two tasks.

Here we tested a strong prediction of the LTP-based theory of interference, namely that disrupting the LTP processes associated with one learning should not only abolish that learning but should release future learning from anterograde interference. We applied continuous theta burst stimulation (cTBS; Huang et al. 2005) in order to causally interfere with the neural processes in early visual cortex immediately after the end of visual training. cTBS is known to act via mechanisms akin to long-term depression (Huang et al. 2005; Huang et al. 2007; Di Lazzaro et al. 2005; Hsieh et al. 2014), thus allowing us to use it to interfere with LTP. We chose the early visual cortex as a target brain region because this area shows changes after visual training (Schoups et al. 2001; Schwartz, Maquet, and Frith 2002; Li, Piech, and Gilbert 2004; Yotsumoto, Sasaki, et al. 2009; Hua et al. 2010; Sasaki, Nanez, and Watanabe 2010; Bang et al. 2014; Bang et al. 2018; Shibata et al. 2017; Rosenthal et al. 2016), suggesting that the early visual cortex is involved in visual perceptual learning. We designed a perceptual learning task that resulted in substantial anterograde interference such that the first learning suppressed the second learning. To anticipate, we found that applying cTBS to the early visual cortex not only abolished the first learning, but also releases the second learning from the anterograde interference. These findings lend direct causal support to the theory that interference occurs due to competition in LTP processes.

## Results

We tested the LTP-based theory of interference. To do so, we examined whether suppressing LTP processes invoked by training on one task disrupts learning on this task and also releases future learning from anterograde interference. We applied cTBS to either the early visual cortex or vertex (control site) immediately after the offset of a training on a Gabor orientation detection task (**Figure 1**). We examined the effects of cTBS on the learning for the orientation trained immediately before cTBS application, as well as for a second Gabor orientation trained one hour later. Learning was quantified as the percent performance improvement from the pre-to post-training tests.

**Figure 1.**
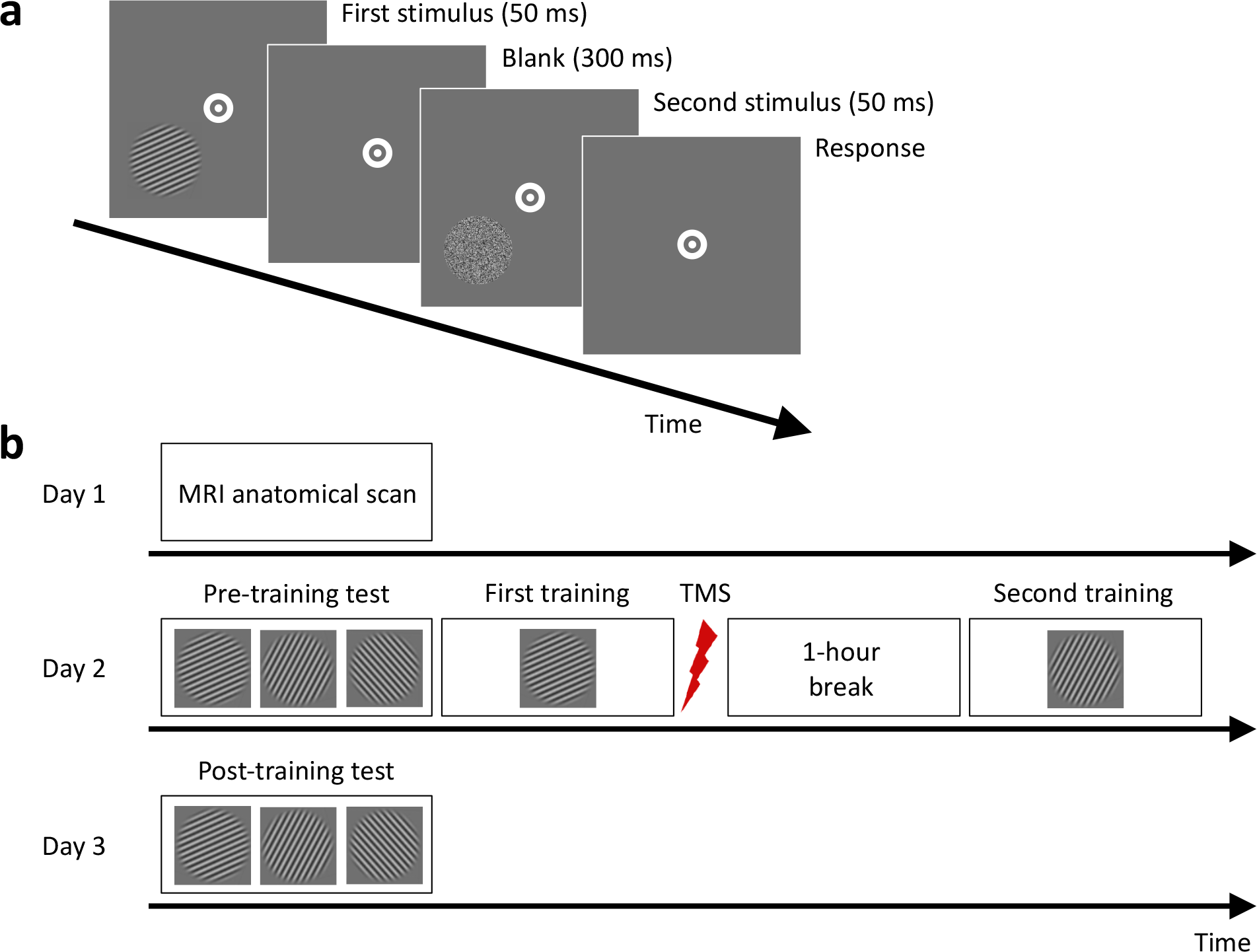
Task and experimental procedure. *(a) Subjects performed a 2-interval forced choice (2IFC) orientation detection task. Two stimuli – a target consisting of a Gabor patch embedded in noise and a non-target consisting of pure noise – were presented in a pre-determined location (either the lower-left or lower-right quadrant) in quick temporal succession. Subjects reported which interval contained the target. (b) The experiment consisted of three days. An MRI anatomical scan was conducted on Day 1 (only for subjects who subsequently received stimulation to their visual cortex). Day 2 began with a baseline pre-training test for each of three stimulus orientations (10°, 70°, and 130°). Subjects were then trained on one randomly chosen orientation and received continuous theta burst stimulation (cTBS) within approximately 2-3 minutes from the training offset. After a one-hour break, subjects completed a second training on a different, randomly chosen orientation. On Day 3, a post-training test assessed how much learning took place for each orientation (trained first, trained second, and untrained).*

To examine the effects of cTBS, we conducted a two-way mixed measures ANOVA on the performance improvement scores with factors cTBS site (visual cortex vs. vertex) and stimulus training (trained first vs. trained second). We found a significant interaction between stimulus training and cTBS site (F(1,23)=11.52, P=0.002; **Figure 2**), demonstrating that cTBS altered the pattern of learning on our task.

**Figure 2.**
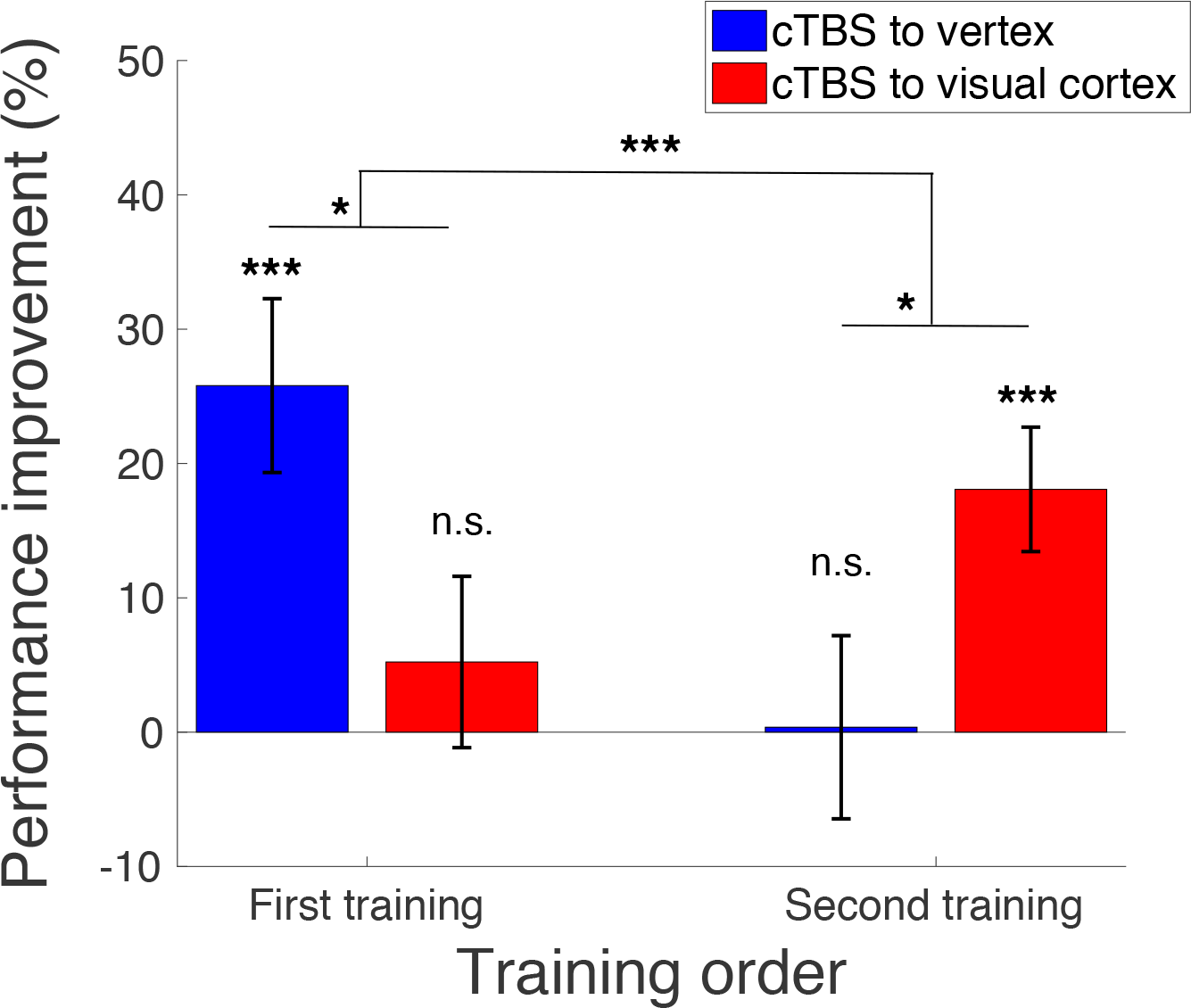
Learning as a function of training order and cTBS site. *We observed an interaction between cTBS site and learning amount (quantified as percent performance improvement) such that cTBS to the visual cortex reduced learning for the first training but increased it for the second training. In other words, cTBS to the visual cortex abolished the pre-stimulation learning and released the second learning from anterograde interference. Vertex served as a control site. Error bars represent s.e.m. * P<0.05, *** P<0.005.*

To understand the exact nature of the cTBS effects, we compared the learning induced by each of the two training periods. We found a significant difference in learning on the first training between subjects who received cTBS on the visual cortex vs. vertex (t(23)=2.26, P=0.03, independent sample t-test), such that there was significant performance improvement after cTBS to the vertex (average improvement = 25.8%; t(11)=3.99, P=0.002, one-sample t-test) but not after cTBS to the visual cortex (average improvement = 5.2%; t(12)=0.82, P=0.43, one-sample t-test). In other words, cTBS to the visual cortex delivered immediately after the end of the first visual training abolished the learning associated with this training.

Next, we performed a similar comparison for the second training and observed the opposite set of results compared to the first training. We again found a significant difference between cTBS to the visual cortex vs. vertex (t(23)=-2.18, P=0.04, independent sample t-test) but this time in the opposite direction: there was no significant learning after cTBS to vertex (average improvement = 0.4%; t(11)=0.06, P=0.96, one-sample t-test) but there was now significant learning after cTBS to the visual cortex (average improvement = 18.1%; t(12)=3.9, P=0.002, one-sample t-test). In other words, by abolishing the learning on the first training, cTBS to the visual cortex released the second training from inhibition thus lending strong support for the notion that interference occurs due to competition of LTP processes.

Finally, we confirmed that our training effects were selective by examining the performance improvement for the untrained Gabor orientation. We found no significant learning in either the subjects who received cTBS to vertex (average improvement = 4.4%; t(11)=0.56, P=0.59, one-sample t-test) or the ones who received cTBS to the visual cortex (average improvement = -3.6%; t(12)=-0.26, P=0.8, one-sample t-test). In addition, there was no significant difference between the learning amount in these two groups for the untrained orientation (t(23)=0.49, P=0.63, independent sample t-test; **Figure 3**).

**Figure 3.**
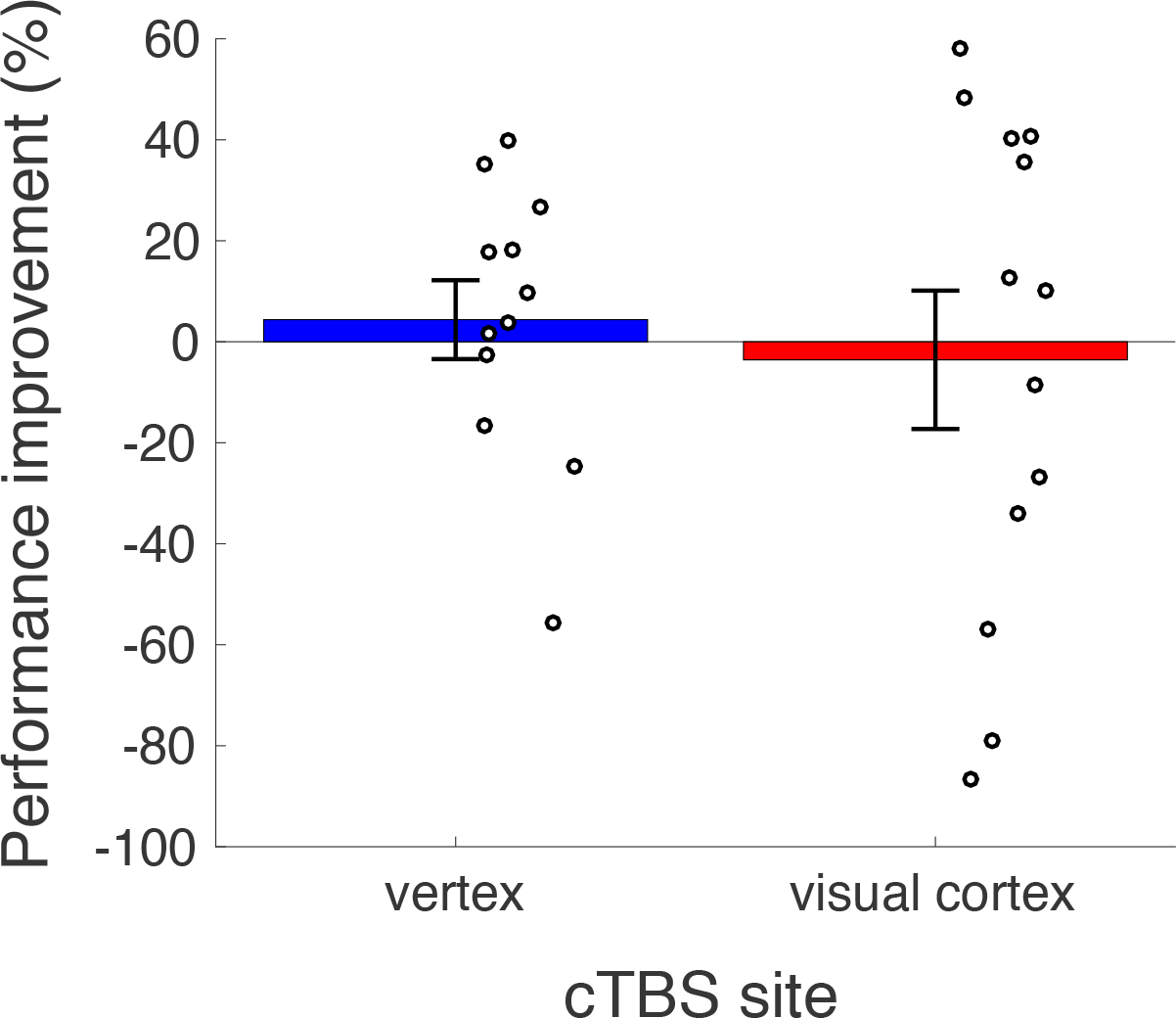
Learning for the untrained orientation. *Subjects did not exhibit any learning (as quantified by percent performance improvement) on the untrained orientation either in the vertex or the visual cortex cTBS conditions. Circles represent individual subjects, error bars represent s.e.m.*

## Discussion

Training two related skills in close succession is known to result in interference (Cantarero et al. 2013; Leow et al. 2013; Wigmore, Tong, and Flanagan 2002; Sing and Smith 2010; Brashers-Krug, Shadmehr, and Bizzi 1996; Shadmehr and Brashers-Krug 1997; Seitz et al. 2005; Yotsumoto, Chang, et al. 2009; Shibata et al. 2017). It is thought that such interference is due to competition in LTP processes associated with each task (Cantarero et al. 2013; Ziemann et al. 2004; Stefan et al. 2006) but direct evidence for this mechanism has been lacking. Here we tested a strong prediction of this LTP-based theory of interference, namely that suppressing LTP processes related to one task should not only disrupt learning on this task but should release future learning from anterograde interference. We applied cTBS to the visual cortex or vertex of human subjects within a few minutes of the training offset. Our results showed that cTBS to the visual cortex abolished performance improvement associated with the training immediately beforehand and released a second training (performed one hour later) from inhibition. These findings provide strong support for the theory that learning interference is based on the competition between LTP-related processes associated with each training.

The notion that new learning leads to LTP has received support in both animal and human subjects. For example, research in rats has demonstrated the existence of learning-induced LTP that leads to a reduced capacity to induce more LTP (Hodgson et al. 2005; Monfils and Teskey 2004; M.-S. Rioult-Pedotti, Donoghue, and Dunaevsky 2007; M. S. Rioult-Pedotti, Friedman, and Donoghue 2000; Mengia -S. Rioult-Pedotti et al. 1998; Sanes and Donoghue 2000). In human subjects learning has been shown to reduce the capacity to induce additional LTP-like plasticity through either the application of precisely-timed pairs of TMS pulses (Ziemann et al. 2004; Stefan et al. 2006; Rosenkranz, Kacar, and Rothwell 2007) or transcranial direct current stimulation (Cantarero et al. 2013). It is widely thought that these learning-induced LTP processes play a critical role in consolidating new skills (Beste et al. 2011; Nicoll 2017; Lynch 2004; Sanes and Donoghue 2000).

Nevertheless, much less is known about the functional role of LTP in the interference between two different episodes of skill learning. It is often thought that LTP processes related to each training compete and it is this competition that leads to interference (Cantarero et al. 2013; Ziemann et al. 2004; Stefan et al. 2006). However, to date there had been no direct evidence for such a mechanism. By causally manipulating the LTP processes related to the first learning, the current study provides causal evidence for such LTP-based theory of interference.

We applied cTBS in order to interfere with LTP neural processes in the visual cortex. Previous studies have already demonstrated that cTBS leads to neural inhibition via mechanisms akin to long-term depression (Huang et al. 2007). For example, within the context of targeting the motor cortex, cTBS suppresses the magnitude of the motor evoked potentials in both humans (Huang et al. 2005; Di Lazzaro et al. 2005) and rats (Hsieh et al. 2014). Therefore, by inducing long-term depression, cTBS is expected to interfere with LTP.

Beyond elucidating the processes behind learning interference, our study also provides causal evidence for the role of the post-training period in the consolidation of new visual skills. Within the domain of motor learning, both studies using behavioral methods and TMS have already established the critical role of the post-training period to skill consolidation. Early research showed that interfering with post-training neural process by introducing a new motor task led to disruption of learning (Brashers-Krug, Shadmehr, and Bizzi 1996; Shadmehr and Brashers-Krug 1997). Direct evidence for the causal role of the immediate post-training period for learning consolidation has been provided by several studies that applied TMS to the primary motor cortex shortly after training on a simple ballistic finger movement task (Baraduc et al. 2004; Muellbacher et al. 2002) or sequence learning task (Robertson, Press, and Pascual-Leone 2005). In all cases, TMS abolished the learning on the motor task. This effect was not observed when TMS was delivered to control brain areas or when TMS was applied to the motor cortex six hours after the practice (Muellbacher et al. 2002). These TMS studies demonstrate that the post-training neural processes have a causal effect in the consolidation of motor learning.

Despite the extensive research in motor learning, evidence for the role of the post-training in consolidating of visual skills is based primarily on behavioral studies (Seitz et al. 2005; Yotsumoto, Chang, et al. 2009; Shibata et al. 2017). A number of previous studies used TMS to understand the mechanisms behind visual perceptual learning (van de Ven and Sack 2013).

Several studies showed that perceptual learning weakens the interference effects of TMS (Corthout et al. 2000; Neary, Anand, and Hotson 2005). Other research demonstrated that TMS delivered during the learning phase of a task leads to interference with long-term memory formation (Giovannelli et al. 2010; Brascamp et al. 2010). Finally, more recent studies used TMS to examine the role of different cortical areas in performing the task after extensive visual training and showed changes in the specialization of cortical areas (Baldassarre et al. 2016; Chen et al. 2016). Although these studies elucidated a number of factors related to visual perceptual learning, none of them examined the role of the immediate post-training period. In fact, only one previous study (De Weerd et al. 2012) applied TMS in the period after the offset of training in order to interfere with visual perceptual learning. In that study two types of stimuli were trained with no break in-between and TMS was applied 45 minutes after the offset of the second training. TMS interfered with learning of the stimulus presented in the targeted retinotopic location but only when it was trained first. Thus, our study is the first to show that TMS applied after a single training can still abolish visual learning and the first to demonstrate a critical role for the period immediately after training offset (since we delivered TMS ∼2-3 minutes rather than ∼45 minutes after training offset) in the consolidation of visual perceptual learning.

An important unresolved question is when competition between LTP processes associated with two tasks would result in anterograde vs. retrograde interference. In our experiment, only anterograde interference was observed in the control condition (cTBS to the vertex). This anterograde interference occurred after a single day of training on two tasks separated by one hour. Previous research suggests that when two episodes of visual training are presented with no delay, then both retrograde and anterograde interference take place (Yotsumoto, Chang, et al. 2009). On the other hand, after five days of training, retrograde interference was observed when the two trainings were presented without delay but not when they were separated by one hour (Seitz et al. 2005). Finally, recent research shows that within a single day of training, the amount of training on the first task determines the extent of retrograde and anterograde interference in the presence of a 30-minute delay (Shibata et al. 2017). These studies differed in the nature of the trained visual stimulus, the amount of training provided per session, the duration of the between-training gap, and the number of sessions trained. Future research should systematically vary these parameters in order to pinpoint when each type of visual learning interference occurs and link these parameters to the properties of LTP.

In summary, our study demonstrates that cTBS disruption of the immediate post-training LTP processes abolishes visual learning and that this abolishment releases later learning from interference. These results provide causal evidence for the theory that learning interference is based on competition between LTP-related processes associated with each learning. Further, taken together with previous research (Baraduc et al. 2004; Muellbacher et al. 2002; Robertson, Press, and Pascual-Leone 2005), our findings establish TMS as a technique that can be used to abolish newly formed memories across a variety of domains and raise the possibility of future therapeutic applications.

## Methods

### Participants

Twenty-five healthy subjects (18 to 25 years old, 12 females) with normal or corrected-to-normal vision participated in this study. Subjects were screened for a history of neurological and psychiatric disorders, as well as for contraindications to TMS and MRI. The study was approved by the Institutional Review Board of Georgia Institute of Technology. All subjects provided written informed consent.

### Procedures

The study consisted of three days (**Figure 1**). On Day 1, subjects who subsequently received stimulation to their visual cortex (N=13) participated in an MRI session. The purpose of the session was to enable us to identify the precise location of stimulation within the early visual cortex. Subjects who received stimulation to a control site (N=12) did not participate in the MRI session. On Day 2, all subjects completed three pre-training tests and two different training periods on a 2-interval forced choice orientation detection task (**Figure 1a**). The three pre-training tests (one block each) were performed for different stimulus orientations (10°, 70°, and 130°), while the trainings (ten blocks each) were performed for two of the three orientations chosen randomly. Immediately after the offset of the first training, subjects received transcranial magnetic stimulation on either early visual cortex or vertex (**Figure 1b**). Subjects were given a one-hour break between the two trainings during which they were asked to watch a nature documentary. On Day 3, subjects completed a post-training test for the same three orientations. Days 1 and 2 were separated by multiple days, while Days 2 and 3 were consecutive.

### Task

Subjects performed a 2-interval forced choice (2IFC) orientation detection task. Each trial began with a 500-ms fixation period. After the fixation period, two stimuli – target and non-target – were presented for 50 ms each, separated by a 300-ms blank period. Subjects indicated which of the two stimuli contained the target (a Gabor patch) by pressing a button on a keyboard. No feedback was provided after the response. For each subject, the stimulus location was pseudo-randomly chosen to be either in the lower-left or lower-right quadrant. The center of the stimulus was placed 4° of visual angle away from the center of the screen in a direction of 45° from vertical toward either lower left or lower right. Once the stimulus location was determined for each subject, all visual stimuli were presented only within that quadrant across the whole experiment. The target was a Gabor patch (diameter = 5°, contrast = 100%, spatial frequency = 1 cycle/degree, SD of Gaussian filter = 2.5°, random spatial phase). We varied the signal-to-noise (S/N) ratio by substituting a random selection of pixels from the Gabor patch with noise pixels (Shibata et al. 2011; Shibata et al. 2017; Seitz, Kim, and Watanabe 2009). The S/N ratio was defined as the percent of pixels that came from the original Gabor patch. The non-target stimulus consisted of pure noise (0% S/N ratio). The target interval (first or second) was determined randomly on each trial. During the entire orientation detection task, subjects were required to fixate on a white bullseye (0.75° radius) at the center of the screen.

The Gabor patches had three possible orientations: 10°, 70°, and 130°. All subjects were tested on all three orientations and received training on two of them. We randomly chose which of the three orientations will be used for the first training, the second training, and which orientation will be untrained.

### Pre-and post-training tests

The pre- and post-training tests were conducted in order to assess the amount of learning that took place. Each pre- and post-training test consisted of three blocks, one for each of the three orientations (10°, 70°, and 130°). The order in which the three orientations were presented during the pre- and post-training tests was determined randomly for each test.

In each pre- and post-training test, we determined the subject-specific threshold intensity for one specific orientation. To do so, we employed a 2-down-1-up staircase procedure within each block. The staircase continuously adjusted the difficulty of the task by altering the S/N ratio.

Each block began with 25% S/N ratio and terminated after 10 reversals. This procedure resulted in subjects completing approximately 40 trials per block. The same staircasing procedure was used during the training blocks as well.

### Analyses

As commonly done for this type of task (Bang et al. 2018; Shibata et al. 2017), we calculated the threshold S/N ratio for each orientation by computing the geometric mean of the S/N ratio during the last six reversals in a block. Based on the obtained intensity thresholds, we computed the performance improvement from the pre- to the post-training test. For each subject and each orientation, the performance improvement was defined as the percent change in the threshold between pre- and post-training tests:

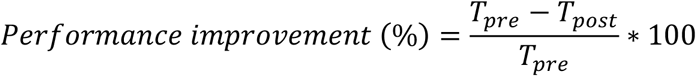
where *T*_*pre*_ and *T*_*post*_ refer to the threshold S/N ratios before and after training. Note that lower threshold values indicate better performance. Statistical tests were performed by conducting two-way mixed measures ANOVAs and following up with independent sample and one sample t-tests.

### Transcranial Magnetic Stimulation

Transcranial magnetic stimulation (TMS) was delivered using a figure-of-eight magnetic coil (MCF-B65) connected to MagVenture MagPro X100 Magnetic Stimulator. We employed continuous theta-burst stimulation (cTBS; Huang et al. 2005), which consists of a series 3-pulse bursts at 50 Hz repeated every 200 ms for 40 seconds (for a total of 600 pulses). Previous studies have shown that the effects of cTBS last for up to an hour after stimulation (Huang et al. 2005).

To calibrate the cTBS intensity, we determined the subject-specific motor threshold on Day 2 shortly before starting the main experiment using established procedures from our laboratory (Rahnev et al. 2013; Rahnev et al. 2016; Rahnev et al. 2012; Shekhar and Rahnev 2018). Briefly, we first applied supra-threshold single pulses around the putative location of the motor cortex in order to identify the best spot for stimulation (defined as the spot that produced maximal finger twitches). We then determined the motor threshold on this location as the minimal TMS intensity required to evoke a visual hand twitch on 5 of 10 consecutive trials. cTBS was then delivered at 80% of the subject-specific motor threshold.

For each subject, we targeted either the vertex or early visual cortex. The vertex served as a control site, whereas the early visual cortex was selected because previous studies suggest that visual learning induces changes in that part of cortex (Schoups et al. 2001; Schwartz, Maquet, and Frith 2002; Li, Piech, and Gilbert 2004; Yotsumoto, Sasaki, et al. 2009; Hua et al. 2010; Sasaki, Nanez, and Watanabe 2010; Bang et al. 2014; Bang et al. 2018; Shibata et al. 2017; Rosenthal et al. 2016). In cases where we stimulated the vertex, we positioned the TMS coil over Cz with the handle extending posteriorly. In cases where we stimulated the early visual cortex, the coil was positioned over the hot spot marked on an anatomical scan of the subject’s brain with the handle extending dorsally. The hot spot was localized in the calcarine sulcus based on anatomical landmarks so as to correspond to the trained region of early visual cortex. Subjects who were trained on the lower left (right) quadrant were stimulated on the corresponding location in the right (left) hemisphere’s dorsal region of the early visual cortex. Stimulation was delivered within 2-3 minutes after the offset of the first training.

### Apparatus

All stimuli were generated in MATLAB using the Psychtoolbox 3 (Brainard 1997). The visual stimuli were presented on an LCD display (1024 A 768 resolution, 60 Hz refresh rate, Mac OS X) in a quiet, dimly-lit room.

### Data Availability

All raw data and analysis codes are freely available at: https://github.com/DobyRahnev/visual_learning_interference_cTBS.

## Acknowledgements

This work was funded by a startup grant to D.R. from the Georgia Institute of Technology.

